# Non-medical prescribing policy in the United Kingdom National Health Service: systematic review and narrative synthesis

**DOI:** 10.1101/582718

**Authors:** Emma Graham-Clarke, Alison Rushton, Timothy Noblet, John Marriott

**Author notes:** Corresponding author (EGC).

## Abstract

Non-medical prescribing was introduced into the United Kingdom (UK) to improve patient care, through extending healthcare professionals’ roles. More recent government health service policy focuses on the increased demand and the need for efficiency. This systematic review aimed to describe any changes in government policy position and the role that non-medical prescribing plays in healthcare provision.

The systematic review and narrative analysis included policy and consultation documents that describe independent non-medical prescribing. A pre-defined protocol was registered with PROSPERO (CRD42015019786). Professional body websites, other relevant websites and the following databases were searched to identify relevant papers: HMIC, Lexis Nexis, UK Government Web Archive, UKOP, UK Parliamentary Papers and Web of Science. Papers published between 2006 and February 2018 were included.

Following exclusions, 45 papers were selected for review; 23 relating to policy or strategy and 22 to consultations. Of the former, 13/23 were published 2006-2010 and the remainder since 2013. Two main themes are identified: chronological aspects and healthcare provision. The impact of government change and associated major healthcare service reorganisation resulted in the publication gap for policy documents. The role of non-medical prescribing has evolved to support efficient service delivery, and cost reduction. For many professions, prescribing appears embedded into practice; however, pharmacy continues to produce policy documents, suggesting that prescribing is not yet perceived as normal practice.

Prescribing appears to be more easily adopted into practice where it can form part of the overall care of the patient. Where new roles are required to be established, then prescribing takes longer to be universally adopted. While this research concerns policy and practice in the UK, this aspect of role adoption has wider potential implications.

## Introduction

Traditionally, prescribing of human medicines had been perceived as a medical role, with only medical professionals and dentists having full prescribing rights in the United Kingdom (UK). Two seminal reports challenged this view; the Cumberlege report [1] which paved the way for limited prescribing by health visitors and district nurses, and the Crown report [2], which recommended extending prescribing rights for the benefit of patients and to utilise the skills of healthcare professionals. The main UK healthcare provider, within which prescribers practice, is the National Health Service (NHS); established in 1948 to provide comprehensive healthcare to all, free at the point of delivery [3]. The UK also has a parallel smaller privately funded healthcare sector. Healthcare policy is directed by the UK government, reflecting the principles of the governing party at the time. Since 1948, this has been one of two main political parties (Labour, Conservative), apart from 2010-2015 when a Conservative and Liberal Democrat coalition was in power. As a general principle, Conservative governments tend to support free markets and expansion of the private sector, whereas Labour governments support the NHS over the private sector. Rising costs and changes in healthcare practice have led to numerous reforms since the NHS was founded but, irrespective of the political stance, the founding principles remain [3, 4].

In 2000 the governing Labour Party published a White Paper ‘The NHS Plan’, which described the government’s intention to modernise healthcare services, breaking down the traditional demarcations between professions and introducing new ways of working to increase healthcare capacity, shorten waiting times, and thus improve the patient experience [5]. Nurse prescribing was highlighted as one of the 10 key roles defined by the Chief Nursing Officer and the White Paper also included broad reference to therapists extending their roles, with prescribing included within this [5]. To support these sweeping changes to traditional practice the government established the Modernisation Agency, tasked with supporting service redesign at a local level [6], and launched a consultation on extending nurse prescribing [7]. This was followed in 2002 by a consultation on the introduction of supplementary prescribing for nurses and pharmacists [8], with approval granted later that year [9].

Supplementary prescribing is described as a voluntary partnership between the supplementary prescriber, the doctor looking after the patient, and the patient. Additionally, a supplementary prescriber can only prescribe medication listed in an agreed clinical management plan [10]. The first supplementary nurse prescribers qualified in 2003, with pharmacists following in 2004. It quickly became apparent that supplementary prescribing, whilst ideal for complex and long-term conditions, had significant limitations with regard to acute care, hampering the government’s desire to enhance patient care through expanding nurse and pharmacist roles and hence improving access to medication. This was articulated clearly in the consultation documents launched in 2005 to investigate expansion into independent prescribing [11, 12]. Legislation to implement independent prescribing by nurses and pharmacists was enacted in 2006 [13], and since that time independent prescribing rights have been gradually extended to a range of healthcare professionals, most recently paramedics [14].

Non-medical prescribing (NMP) is the umbrella term used to cover prescribing by professions other than doctors. The initial focus of government policy with regard to NMP was the desire to improve patient access to medicines. However, more recent documents from NHS England have focused on the increased demand for services and the need to drive efficiency so that maximum benefit can be obtained from the limited NHS budget [15, 16]. The role of NMP has been less apparent in these later documents, and it is unclear if this reflects a change in government policy.

The present study aimed to conduct a systematic literature review investigating potential changes in UK Government policy position with regard to NMP, since the introduction of independent prescribing for nurses and pharmacists. The study also aimed to determine the current role of independent NMP in the delivery of healthcare in the NHS.

## Method

### Protocol and registration

A systematic review with narrative synthesis was conducted to explore the evolution of government policy concerning independent NMP in the UK. A predefined protocol was developed following the PRISMA-P statement [17] and registered with PROSPERO (CRD42015019786) (S1 Protocol). The results are reported following the PRISMA statement (S1 appendix) [18].

### Eligibility criteria

Documents describing policy concerning independent NMP in the UK were included. These included White and Green papers, policy statements, consultation documents and reports. Documents published since 2006 were included, as the legislation permitting nurse and pharmacist independent prescribing was enacted in that year [13].

### Information sources

Advice was taken from expert librarians regarding appropriate electronic databases and websites to search (listed in **Error! Reference source not found.**) and to aid development of search strategies. Broad search terms (e.g. prescribing, non-medical) were used to capture as wide a range of documents as possible. Boolean operators and truncation were used if the database supported them. Iterative and ‘snowball’ search techniques were employed [19], with the primary searches complete to the end of February 2018, and secondary searches conducted as necessary (S2 Appendix). Documents obtained were mapped to identify gaps (for example, documents relating to the consultation process or profession specific policy documents) enabling targeted secondary searches to be conducted. Relevant citations in the reviewed documents were also obtained and personal files searched [19]. Full texts of the selected documents were screened to remove those that did not meet the eligibility criteria.

**Table 1.**
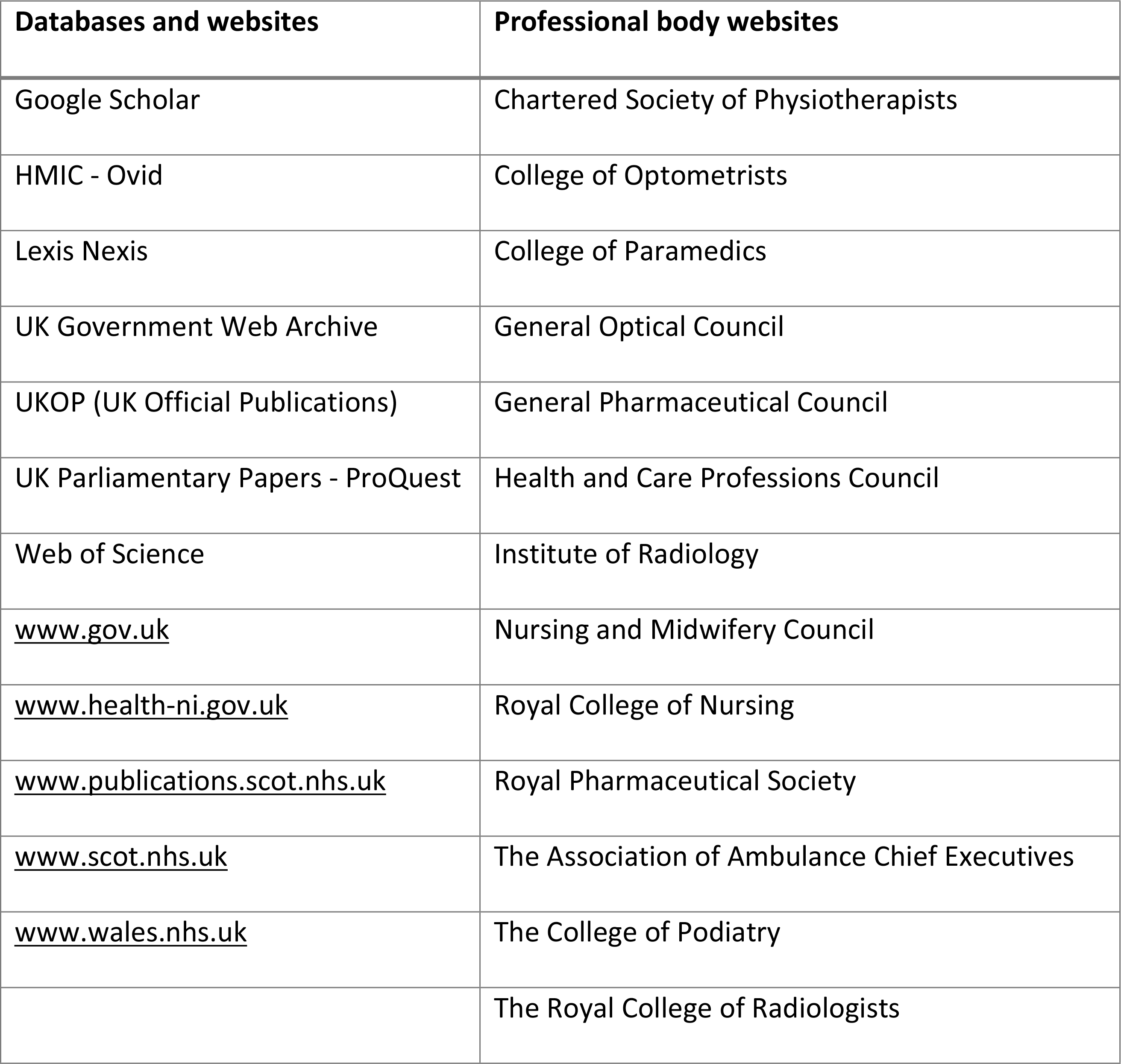
Databases and websites searched

| Databases and websites | Professional body websites |
| --- | --- |
| Google Scholar | Chartered Society of Physiotherapists |
| HMIC - Ovid | College of Optometrists |
| Lexis Nexis | College of Paramedics |
| UK Government Web Archive | General Optical Council |
| UKOP (UK Official Publications) | General Pharmaceutical Council |
| UK Parliamentary Papers - ProQuest | Health and Care Professions Council |
| Web of Science | Institute of Radiology |
| www.gov.uk | Nursing and Midwifery Council |
| www.health-ni.gov.uk | Royal College of Nursing |
| www.publications.scot.nhs.uk | Royal Pharmaceutical Society |
| www.scot.nhs.uk | The Association of Ambulance Chief Executives |
| www.wales.nhs.uk | The College of Podiatry |
|  | The Royal College of Radiologists |

### Study selection

Two reviewers (EGC and TN) independently conducted each stage, resolving differences by discussion, with a third reviewer (AR) available if required for mediation [20]. Numbers excluded were recorded [18, 20].

### Data collection process and data items

Selected papers were entered into a Microsoft^®^ Excel for Mac (version 16) spreadsheet. Home nation and professions covered by the reference were noted, and whether the reference related to policy or consultation. The full texts were read, and notes made of any reference to NMP, including the context.

### Risk of bias assessment

Unlike research papers, whether qualitative or quantitative in nature, policy and consultation documents are not developed according to well-recognised principles. Risk of bias assessment is therefore not appropriate for this type of document and was not conducted. Policy documents are liable to be biased towards the ethos of the government in power at the time and documents produced by profession specific bodies towards their profession. The results will be reported according to the relevant government era and, where appropriate, the specific professional body.

### Data syntheses

Narrative synthesis was conducted on the selected documents [21, 22]. Following tabulation and data extraction, the selected documents were grouped depending on whether they concerned policy or consultation. To aid this process and to visualise the time distribution they were also plotted on a timeline, with a further timeline developed for the consultation documents. Using these techniques, a narrative summary was able to be developed by one researcher (EGC), and the findings were then debated and critically assessed by all authors to reach agreement.

One of the authors (EGC) is a practising pharmacist independent prescriber and NMP lead for an acute trust. In this role they support other non-medical prescribers and have an interest in NMP developments. This researcher standpoint is balanced by the other authors, who do not have prescribing qualifications.

## Results

### Paper selection and characteristics

The search strategy identified 99 full text articles to be assessed for inclusion. Following exclusions, 45 papers were included in the review (Fig 1).

**Fig 1.**
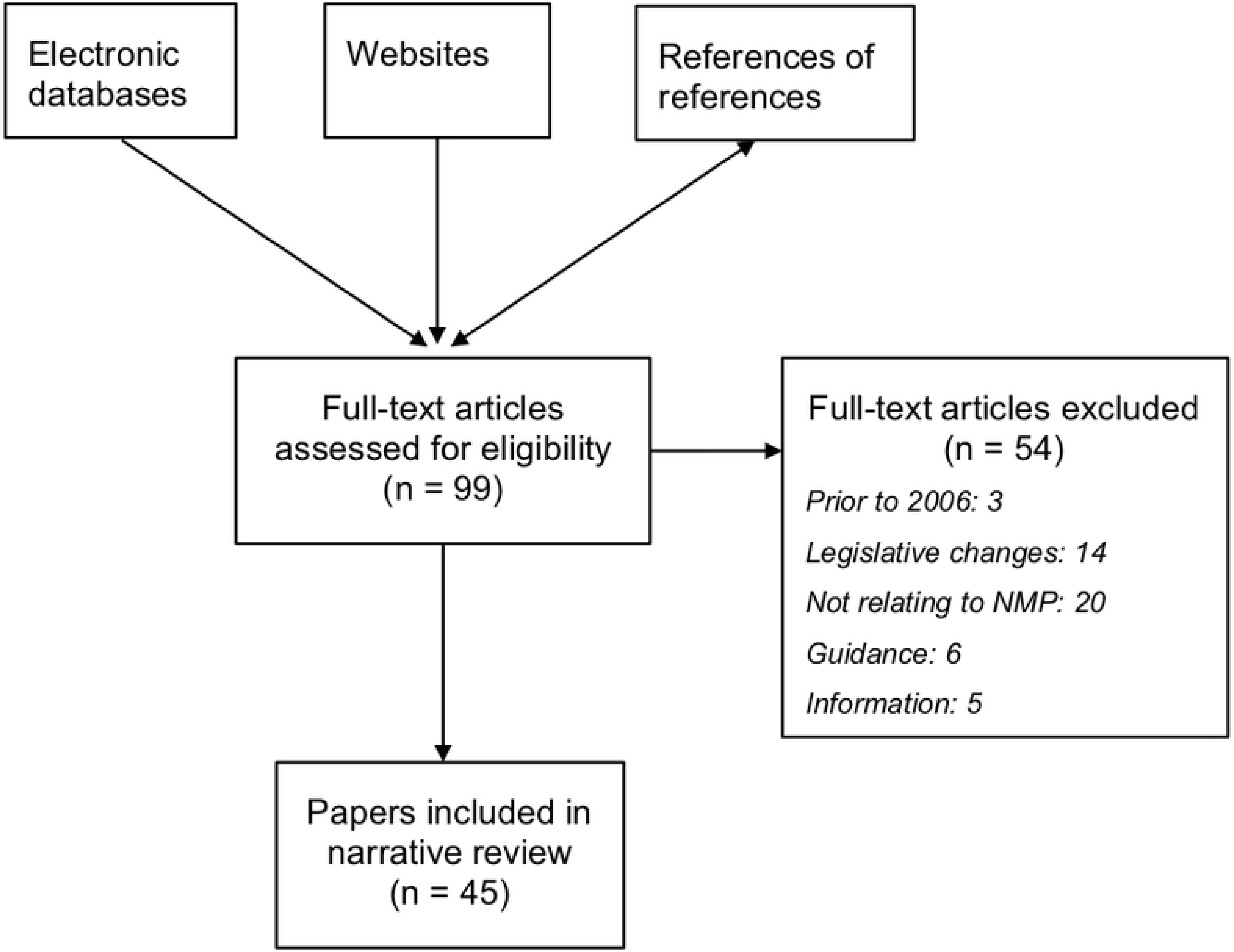
PRISMA paper selection flow diagram.

Of the included papers, 23 relate to policy or strategic report documents (see **Error! Reference source not found.**), and 22 to the consultation process concerning extension of independent NMP responsibilities to various healthcare professions (see **Error! Reference source not found.**).

**Table 2.**
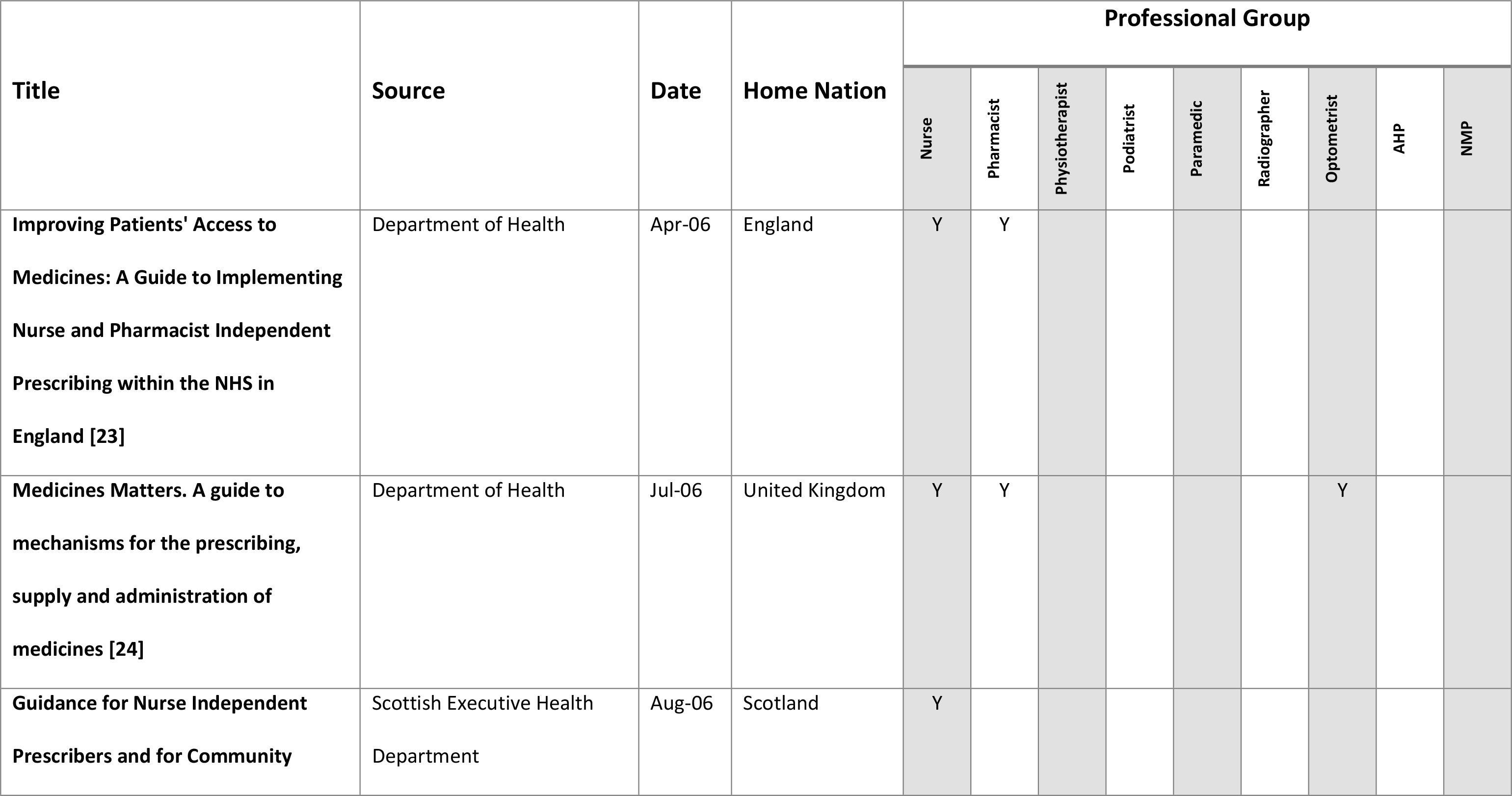
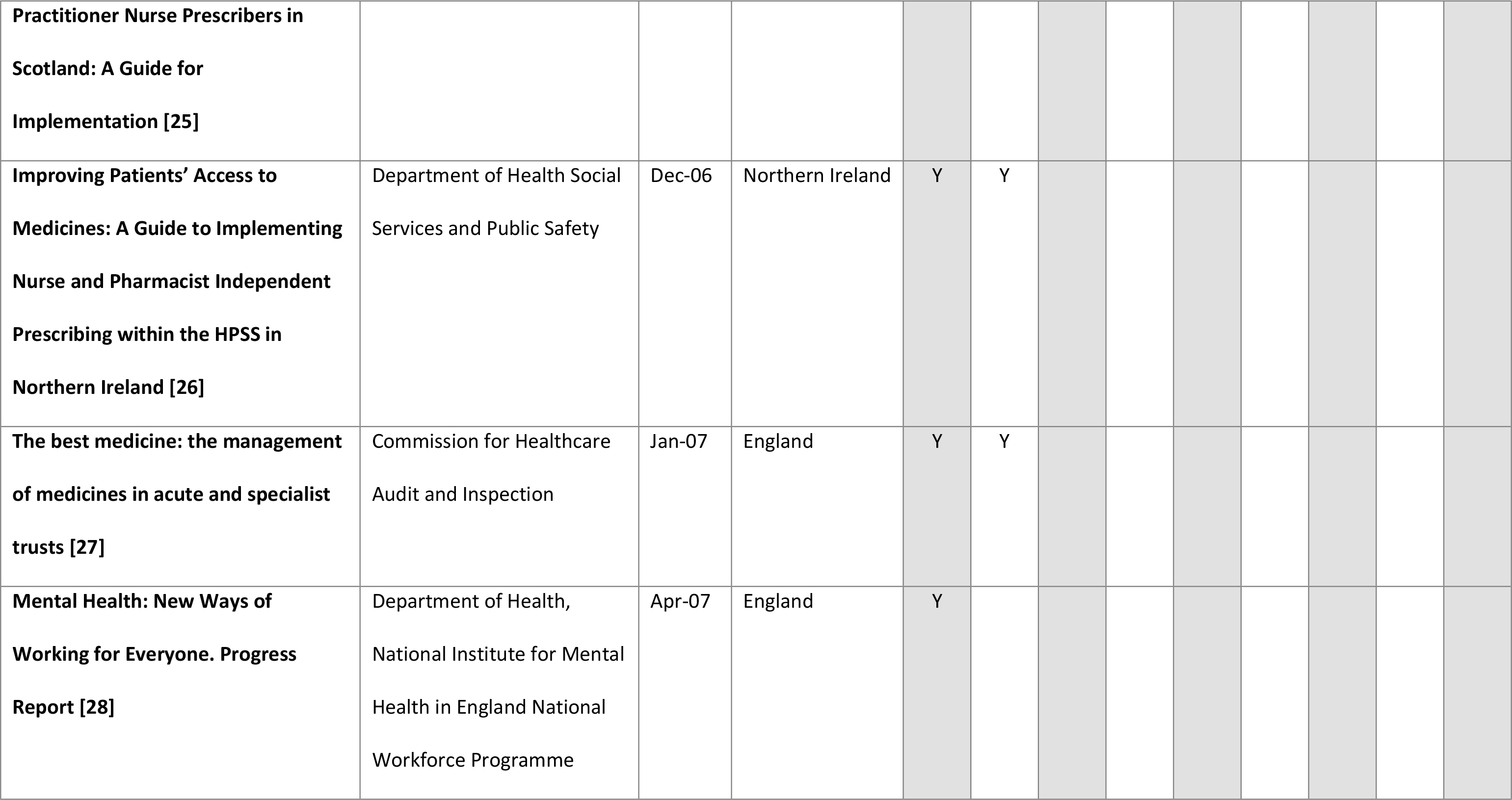
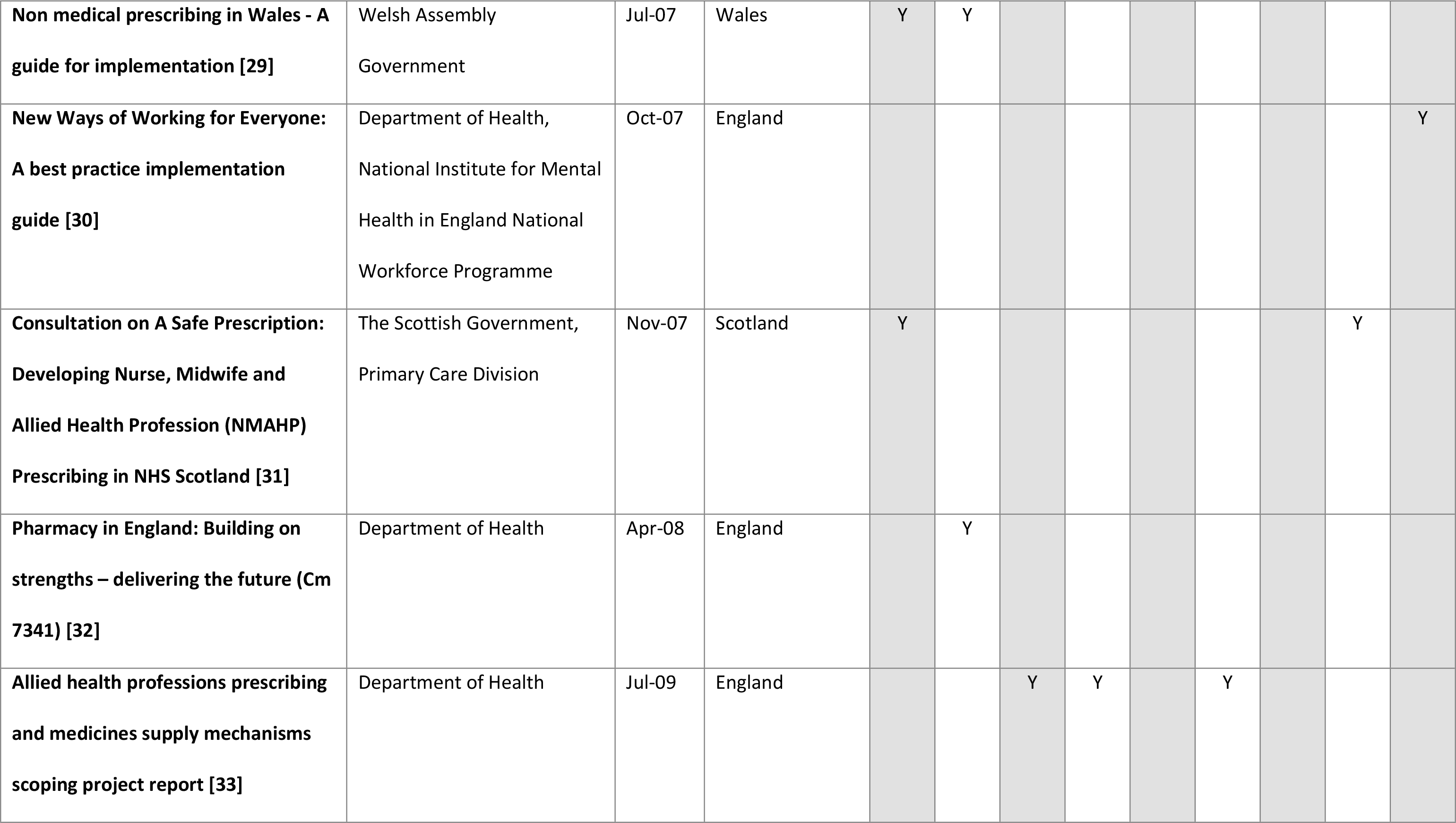
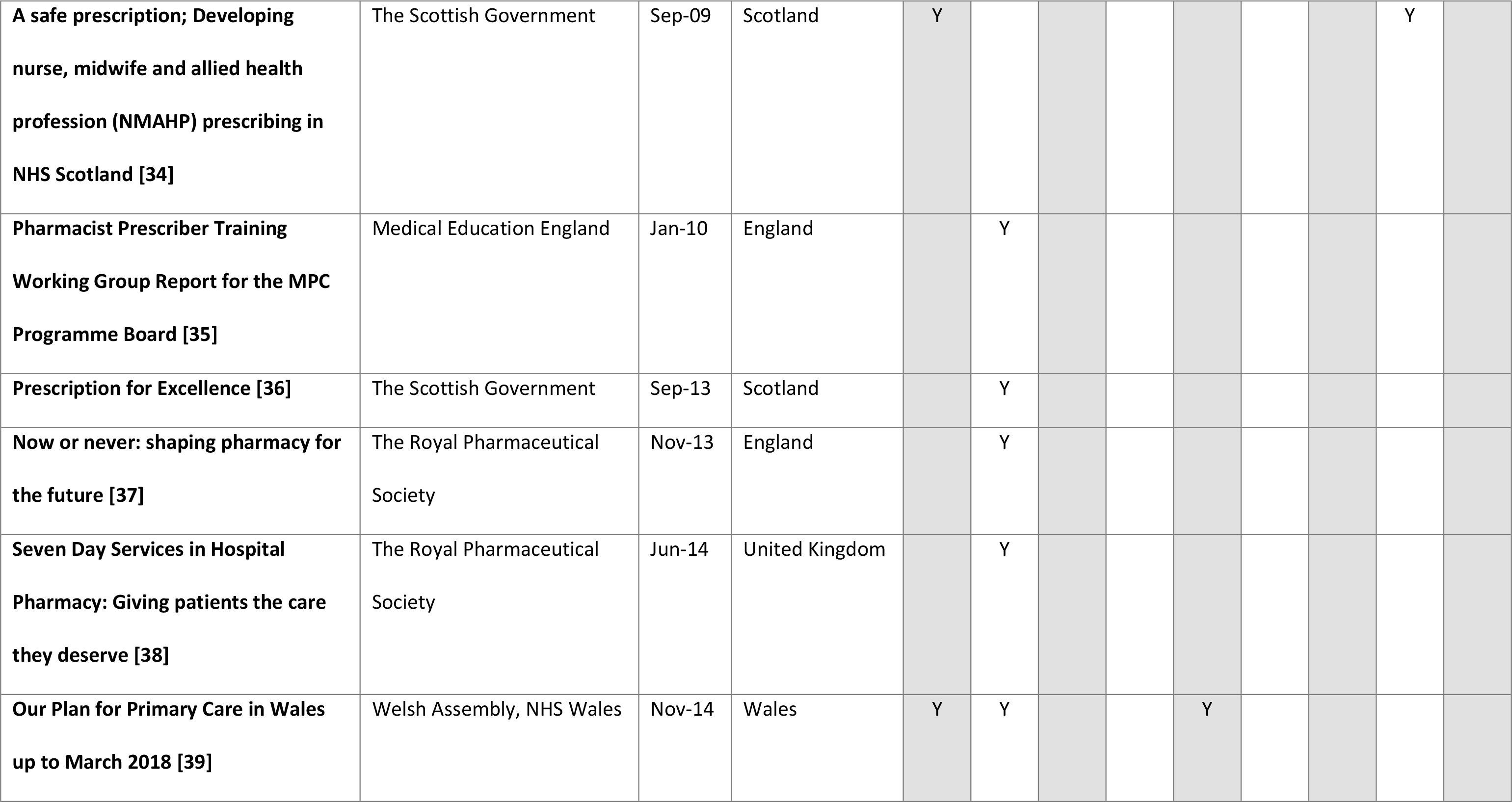
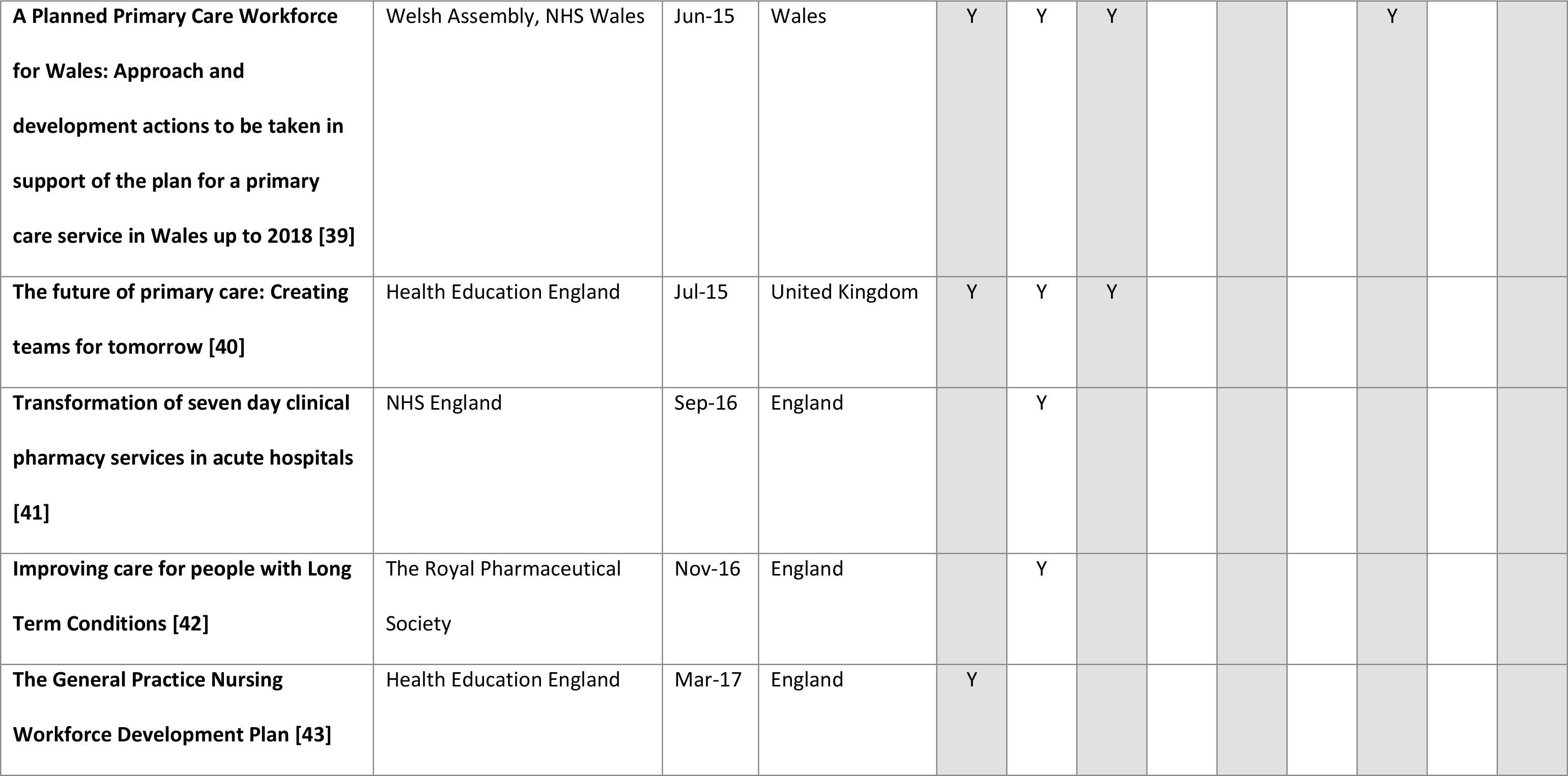
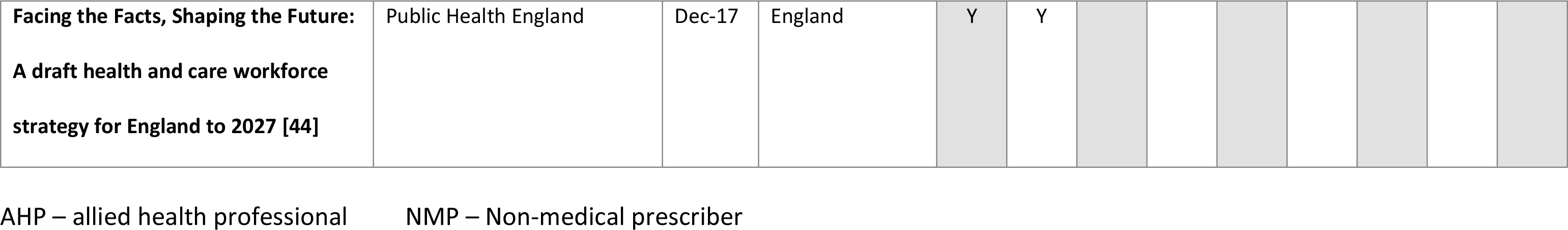
Policy and Strategic Report Documents

**Table 3.** Consultation Documents

| Title | Source | Date | Home Nation | Professional Group |  |  |  |  |  |  |
| --- | --- | --- | --- | --- | --- | --- | --- | --- | --- | --- |
|  |  |  |  | Nurse | Pharmacist | Physiotherapist | Podiatrist | Paramedic | Radiographer | Optometrist |
| **Consultation on proposals to introduce independent**<br>**prescribing by optometrists (MLX 334) [45]** | Medicines &<br>Healthcare<br>Products<br>Regulatory Agency | Aug-06 | United Kingdom |  |  |  |  |  |  | Y |
| **Public consultation - independent prescribing of controlled drugs by nurse and pharmacist independent prescribers (MLX338) [46]** | Home Office, Drug Strategy Unit | Mar-07 | United Kingdom | Y | Y |  |  |  |  |  |
| **Public consultation (MLX 334): Proposals to introduce independent prescribing by optometrists – outcome [47]** | Medicines & Healthcare Products Regulatory Agency | Aug-08 | United Kingdom |  |  |  |  |  |  | Y |
| **Proposals to introduce prescribing responsibilities for paramedics: stakeholder engagement [48]** | Department of Health | Mar-10 | United Kingdom |  |  |  |  | Y |  |  |
| **Engagement exercise: To seek views on possibilities for introducing independent prescribing responsibilities for podiatrists [49]** | Department of Health | Sep-10 | United Kingdom |  |  |  | Y |  |  |  |
| **Engagement exercise: To seek views on possibilities for introducing independent prescribing responsibilities for physiotherapists [50]** | Department of Health | Sep-10 | United Kingdom |  |  | Y |  |  |  |  |
| **Proposals to introduce independent prescribing by podiatrists: impact assessment [51]** | Department of Health | Jul-11 | United Kingdom |  |  |  | Y |  |  |  |
| **Consultation on proposals to introduce independent prescribing by podiatrists [52]** | Department of Health | Sep-11 | United Kingdom |  |  |  | Y |  |  |  |
| **Consultation on proposals to introduce independent prescribing by physiotherapists [53]** | Department of Health | Sep-11 | United Kingdom |  |  | Y |  |  |  |  |
| **Summary of the Commission on Human Medicines meeting held on Thursday 17th & Friday 18th May 2012 [54]** | Commission on Human Medicines | May-12 | United Kingdom |  |  | Y | Y |  |  |  |
| **Summary of Public Consultation on Proposals to Introduce Independent Prescribing by Physiotherapists [55]** | Department of Health | Jul-12 | United Kingdom |  |  | Y |  |  |  |  |
| **Proposals to introduce independent prescribing by physiotherapists: impact assessment [56]** | Department of Health | Jul-12 | United Kingdom |  |  | Y |  |  |  |  |
| **Summary of Public Consultation on Proposals to Introduce Independent Prescribing by Podiatrists [57]** | Department of Health | Jul-12 | United Kingdom |  |  |  | Y |  |  |  |
| **Independent prescribing by radiographers: Impact Assessment [58]** | NHS England | Jan-15 | United Kingdom |  |  |  |  |  | Y |  |
| **Consultation on proposals to introduce independent prescribing by radiographers across the United Kingdom [59]** | NHS England | Feb-15 | United Kingdom |  |  |  |  |  | Y |  |
| **Consultation on proposals to introduce independent prescribing by paramedics across the United Kingdom [60]** | NHS England | Feb-15 | United Kingdom |  |  |  |  | Y |  |  |
| **Proposal to introduce independent prescribing by paramedics: impact assessment [61]** | NHS England | Feb-15 | United Kingdom |  |  |  |  | Y |  |  |
| **Commission on Human Medicines and Expert Advisory Group Final Summary Minutes [62]** | Commission on Human Medicines | Oct-15 | United Kingdom |  |  |  |  | Y | Y |  |
| **Independent prescribing by therapeutic radiographers [63]** | NHS England | Jan-16 | United Kingdom |  |  |  |  |  | Y |  |
| **Summary of the responses to the public consultation on proposals to introduce independent prescribing by paramedics across the United Kingdom [64]** | NHS England | Feb-16 | United Kingdom |  |  |  |  | Y |  |  |
| **Summary of the responses to the public consultation on proposals to introduce independent prescribing by radiographers across the United Kingdom [65]** | NHS England | Feb-16 | United Kingdom |  |  |  |  |  | Y |  |
| Summary of the Commission on Human Medicines meeting<br>held on Thursday 7th September 2017 [66] | Commission on<br>Human Medicines | Sep-17 | United Kingdom |  |  |  |  | Y |  |  |

The policy and strategic report documents relate to a single profession (nursing 3, pharmacy 7), multiple professions (12), or generic NMP (one). The majority concern matters in the home nations (England 12, Scotland 4, Wales 3 and Northern Ireland with only 3 concerning the United Kingdom. They can be divided into two chronological eras, with just over half published between 2006 and 2010, and the remainder published since 2013 (Fig 2).

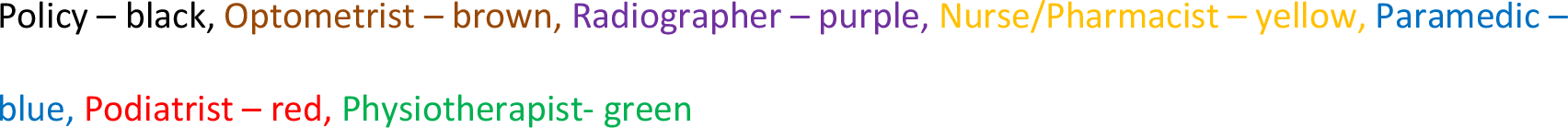

**Fig 2.**
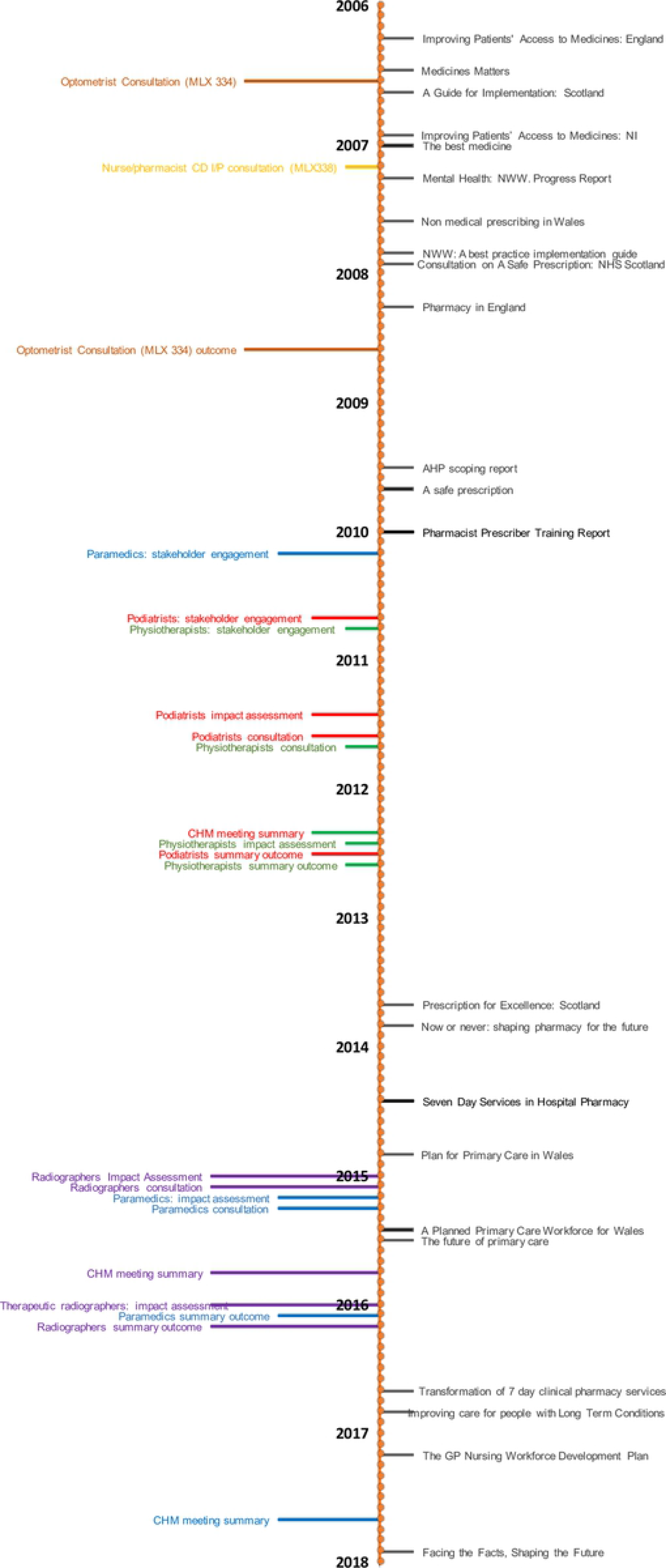
Timeline of selected documents.

### Synthesis of results

#### The Labour Government era 2006-2010

Four of the early documents comprised guidance issued by the home nations to support NMP. These were released as the relevant regulations governing prescribing were amended to permit independent NMP. The first was released by the Department of Health in April 2006, coinciding with the initial changes in legislation and regulations permitting independent prescribing by nurses and pharmacists [13, 23, 67]. This was followed by Scotland’s guidance, released in July 2006, Northern Ireland’s guidance in December 2006 and the Welsh guidance in 2007 [25, 26, 29].

All four documents are similar in nature; however, Scotland’s relates to nurse prescribing only whereas the other three relate to nurse and pharmacist prescribing. This reflects the changes made by the home nations whereby England, Wales and Northern Ireland each introduced nurse and pharmacist independent prescribing simultaneously, whereas Scotland introduced nurse independent prescribing first, followed a year later by pharmacist independent prescribing. Although the bulk of these documents relates to practical implementation guidance, each states the core policy drivers behind NMP which were:

- improving patient care, without reducing safety
- making it easier to patients to access the medicines they require
- increasing patient choice
- utilising the skills of health professionals
- supporting team working

The Welsh guidance included the additional benefits of improving healthcare capacity and enhancing patient access for advice and services.

Scotland conducted a prescribing strategy consultation exercise, with the final strategy launched in 2009 [31, 34]. These documents covered independent prescribing by nurses and midwives and supplementary prescribing by allied health professionals but not pharmacist prescribers. They highlighted the variable adoption of NMP across Scotland and had the aim of improving uptake of NMP to support the NHS boards in delivering patient centred care.

The remaining prescribing specific documents in this era were the scoping report on Allied Health Professional (AHP) prescribing and a report on pharmacist prescribing training [33, 35]. The former reviewed the developing role of AHPs and highlighted some of the limitations resulting from their inability to prescribe; identifying which professions would benefit most from the ability to prescribe, either independently or as a supplementary prescriber, and also which professions should not become prescribers. Additionally, the professions were prioritised, with physiotherapy and podiatry identified as high priorities for independent prescribing, followed by radiology. The latter document reviewed pharmacist prescribing experiences and recommended several changes to training, both at undergraduate level and regarding the prescribing course.

The remaining documents produced in this era, although generic in nature, include references to NMP. The first was a Department of Health document released in 2006 providing further guidance on medicine supply and reiterating the drive behind NMP [24]. The document included several proposed next stages for NMP:

- To consult on optometrist independent prescribing
- To promote nurse and pharmacist independent prescribing
- To review the prescribing needs of emerging roles

This was followed by the Audit Commission report in 2007 on medicines management, which mentioned the development of nurse and pharmacist prescribing and described the distribution of prescribers at that time [27]. Data collection had been in 2005 and 2006 and therefore the majority of these data would have been collected from supplementary prescribers. They recommended that trusts identify where NMP would provide the maximum benefit clinically and that work should be performed to identify why some non-medical prescribers did not prescribe regularly.

The “New Ways of Working in Mental Health” project released two documents in 2007, a progress report and an implementation guide [28, 30]. The progress report reiterated the five core drivers behind NMP and described how NMP should be incorporated into the changes in working practice such as multidisciplinary team working. The implementation guide provided theoretical examples of changed practice which incorporated NMP.

The final document in this era was the pharmacy White Paper [32]. This highlighted the roles that pharmacists could play in improving the healthcare of patients, including the example of prescribing in long-term conditions. Although some case studies were described, most of the suggested roles for prescribers were aspirational.

#### The Coalition and Conservative Governments era 2013-2017

The first two documents in this era both concerned the role of pharmacy in providing patient centred health care. The first of these was the Scottish Government’s vision for pharmacy which envisaged integration of pharmacists into all aspects of healthcare [36]. Central to this vision was the aim of having all pharmacists qualified as independent prescribers. The second document was a report by the Royal Pharmaceutical Society on pharmacy activity and future potential [37]. Various examples of prescribing practice are described but the comment is made that it is not sufficient simply to provide prescribing courses, that roles must also be developed that utilise this skill. The report contrasts the English and Scottish governments approach to pharmacy, to the detriment of the English government’s approach.

There are three further pharmacy specific documents in this era, with two of these concerning seven-day hospital pharmacy services. The first was a report by the Royal Pharmaceutical Society discussing potential approaches to providing a seven-day service and the associated challenges [38]. Examples where seven-day pharmacy services had been implemented were given, with many of the contributors anticipating the use of pharmacist prescribers to support delivery. The second report, from NHS England, describes the need to deliver clinical pharmacy services seven days a week, highlighting the impact that pharmacy services make and describing the importance of prescribing to support the multi-professional team [41]. The final pharmacy specific document was the Royal Pharmaceutical Society produced policy paper, concerning care for patients with long-term conditions [42]. This highlights the role that pharmacists can play in supporting these patients, and makes a number of key recommendations, the first of which is that pharmacists should have the opportunity to become prescribers.

The Welsh Assembly produced a plan for primary care in 2014, followed by a primary care workforce development plan in 2015 [39, 68]. The first of these documents highlighted the increasing pressure on general practice from a combination of increasing demand, a shortage of general practitioners and financial constraints. The focus was on health rather than ill-health and to provide person centred care within the local community, using the most appropriate healthcare professional for the task. Advanced practice such as NMP was seen to relieve pressure on general practitioners. The associated workforce plan described the potential role of NMP for various professions and provided examples.

A Health Education England commissioned report on primary care, published in 2015, described how primary care could be delivered using a wide range of healthcare professionals [40]. Included in the recommendations was the role of the prescribing pharmacist, and the potential for physiotherapist prescribers. This was followed in 2017 by the general practice nursing workforce plan [43]. Prescribing is described as complementing the nursing role, but challenges are acknowledged particularly in enabling time for training. Finally, there was the draft workforce strategy for England which was released for consultation in December 2017 [44]. This specifically mentioned prescribing in the pharmacy section and also commented that increased numbers of nurse prescribers would be required in the community and primary-care sectors. No mention was made of prescribing by any other non-medical healthcare professional.

#### Consultation documents

Two consultations were launched during the period 2006-2008; the first concerned the introduction of independent prescribing for optometrists, and the second regarding controlled drug prescribing by nurse and pharmacist independent prescribers. The consultation process for the introduction of independent prescribing by optometrists was launched in August 2006, with the outcome announced in 2008, and associated legislation passed the same year [45, 47, 69]. This time period contrasts with the second consultation in 2007 on controlled drug prescribing, where agreement that this should be permitted was reached, but changes in legislation were not enacted until 2012 [46, 70, 71].

Following the 2009 AHP scoping report, stakeholder engagement exercises were launched in 2010 to investigate independent prescribing rights for both podiatry and physiotherapy, followed by consultation exercises in 2011 and the outcome and approval in 2012, the whole process taking a little under two years [49–57]. The consultation for radiographers was launched in 2015 with approval, for therapeutic radiographers only, granted in 2016 (diagnostic radiographers were excluded) [58, 59, 62, 63, 65]. These relatively short consultation exercises contrast strongly with that of the paramedics. The initial document mentioning paramedic prescribing was published in 2005 [72], with the stakeholder engagement exercise held in 2010, a year before that of the podiatrists and physiotherapists [48–50]. The potential for paramedic prescribing was reiterated in the 2013 urgent care report, which described the changing role of paramedics, and the potential for further role extension [73]. The paramedic and radiographer consultation exercises ran simultaneously, but final approval for paramedics was only granted in 2017 [60, 61, 64, 66, 74]. A comment is made in the related paramedic impact assessment that the consultation exercise was delayed because of capacity issues [61]. The relative timescales are visually depicted in Fig 3.

**Fig 3.**
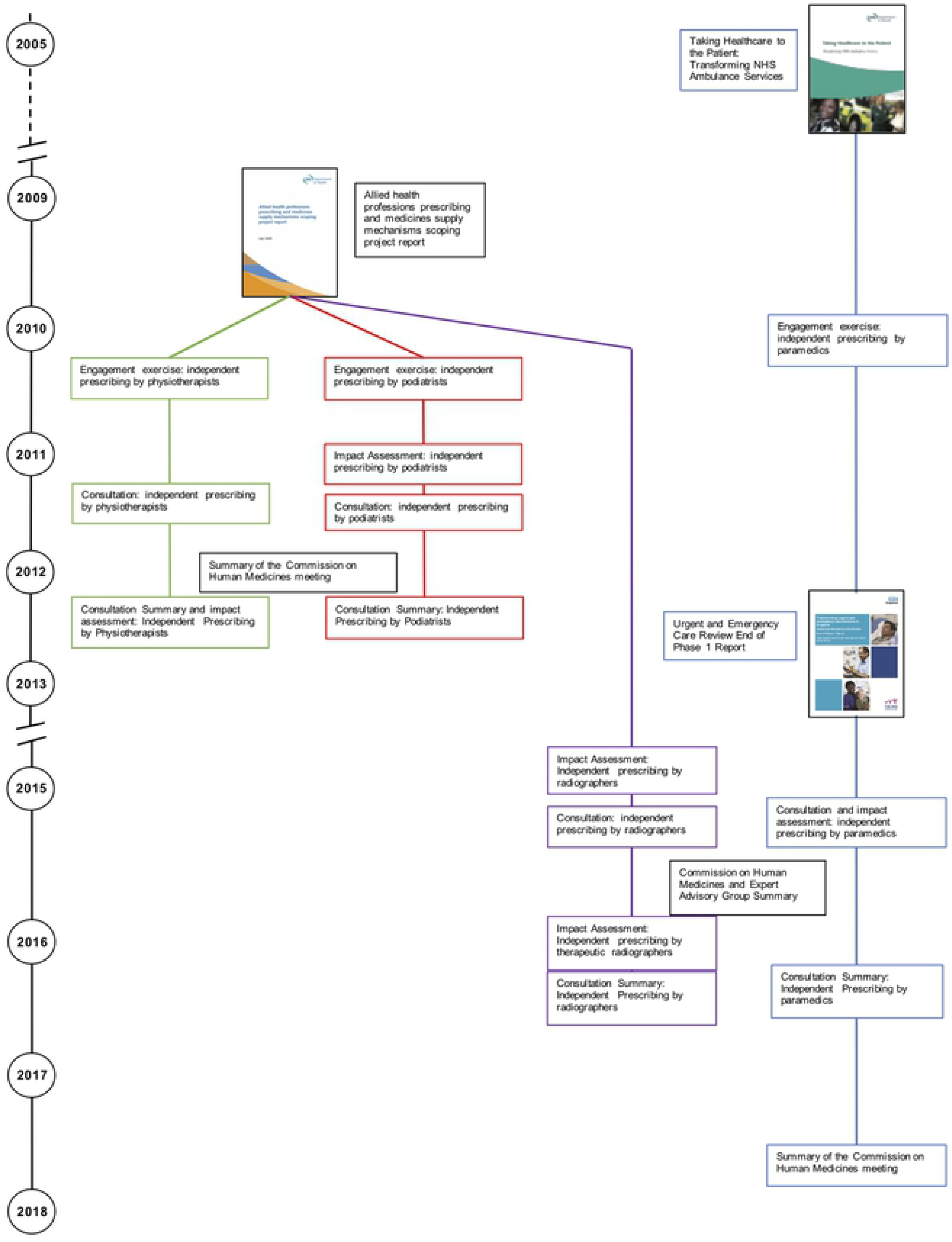
Consultation Timeline.

## Discussion

### Summary of evidence

This is the first such review bringing together the UK policy documents concerning NMP to describe the role of this evolving skill. Review of the evidence reveals two main themes, which are expanded on below. The first theme highlights issues arising from inspecting the chronological aspects of the selected documents. The second theme covers the evolving approach to healthcare provision and describes how NMP has become embedded into routine practice for many non-medical prescribers. However, differences in practice remain and these are highlighted.

#### Chronological aspects

Inspection of the timeline of included papers reveals a noticeable gap between 2010 and 2013, when no reports or strategic documents concerning NMP were released by a government body. The beginning of this period coincides with the change in government in 2010 from Labour to the Coalition. Two factors are likely to be responsible for this dearth of publications. Firstly, the Coalition embarked on an overall reorganisation of the NHS in England, initiated in the 2010 White Paper ‘Equity and Excellence’, and enacted through the Health and Social Care Act in 2012 [75, 76]; focussing on the high level structure rather than finer detail. Secondly, the country had been in economic recession since 2008 and the Coalition’s 2010 budget introduced austerity measures designed to reduce the nation’s budget deficit and improve economic growth [77, 78]. The government attempted to protect the NHS from financial cuts implemented more generally across all services, however the funding growth rate for the NHS in England was curtailed to 1.4% a year compared with 6% a year under the previous Labour government [79]. Government priorities were therefore concerned with major reform of the NHS structure and introduction of commissioning groups, rather than the continued development of existing practices.

The change in government also probably explains the delay in extending controlled drug prescribing for nurses and pharmacist independent prescribers. Extending controlled drug prescribing rights requires the agreement of the Department of Health, the Home Office, the Medicines and Healthcare Products Regulatory Agency and the Advisory Council on the Misuse of Drugs (ACMD), and, subsequently, amendments to the Misuse of Drugs Regulations 2001 and medicines legislation [46]. The consultation closed in June 2007, and in November 2007 the ACMD wrote to the Under-Secretary of State at the Home Office, and the Minister of State for Public Health at the Department of Health, to support the proposals and the change in legislation [80]. However, the required change in legislation was only enacted in 2012, and it can be surmised that with the Coalition’s priorities focused on reorganisation of the whole NHS, extending controlled drug prescribing to nurse and pharmacist independent prescribers was accorded low priority [70, 71].

The consultation processes for the AHPs (physiotherapists, podiatrists and radiographers) were all concluded within a reasonable timeframe, despite the change in government occurring between publication of the AHP scoping report and initiation of the physiotherapy and podiatry consultation exercises [33, 49, 50]. The AHP scoping report had demonstrated a clear role for prescribing for each of these professions in streamlining and improving patient care. In addition, the report prioritised which professions should be considered first, taking into consideration the strength of case for prescribing for each profession and the capacity of the Department of Health and Medicines and Healthcare Products Regulatory Agency to conduct the necessary consultations. As an aside, the consultation exercises reflect the NHS reorganisation, with the physiotherapy and podiatry consultation exercises conducted under the auspices of the Department of Health, and subsequent consultation exercises under NHS England.

In comparison, the lack of clarity concerning how prescribing would be utilised by paramedics, and their evolving role, explains the extended time period between the initial recommendation regarding paramedic independent prescribing and final approval. At the time of the initial report paramedics had recently become registered with the Health Care Professions Council, and the NHS advanced practice role was developing [72]. Consequently, the training focus shifted from resuscitation, to assessing and treating the patient at home. By the time of the urgent care report in 2013, treatment by a paramedic at home was considered an essential component of the strategy to reduce demand on emergency care services [73]. Furthermore, when the formal consultation process began, advanced paramedics had started to work in a range of settings such as emergency care departments as well as the more traditional ambulance service. Following the consultation, the Commission on Human Medicines (CHM) was unable to recommend prescribing by paramedics because of concern that paramedics would need training in a large range of conditions to ensure patient safety [62]. The minutes for the 2017 CHM meeting simply say that they endorse the recommendations for independent prescribing for paramedics, and it is to be presumed that they had been provided with reassurance concerning the training and role of paramedics [66].

#### Healthcare provision - evolution of policy

The five drivers for prescribing documented in the implementation guidance reiterated the aims of the 2000 NHS White Paper to improve patient care and break down the traditional demarcations between professions [5, 23, 25, 26, 29]. These and other early documents such as the “Medicines Matters, and the Mental Health New Ways of Working” project were published before full independent prescribing was embedded [24, 28, 30]. As such, they discuss the potential for NMP to improve patient care and, in particular with the mental health documents, develop new ways of working. Medicines Matters explicitly commented that NMP was unsuitable for patients with complex conditions, recommending the use of supplementary prescribing instead [24]. The pharmacy White Paper listed prescribing as one of the activities that pharmacists could undertake including the care of long-term conditions but many of the examples are theoretical [32]. The 2009 AHP scoping report highlights the changing role of, for example physiotherapists or podiatrists, commenting that they may now be responsible for a full package of patient care but were hampered by the inability to prescribe independently [33]. Again, this document describes potential or theoretical benefits.

However, when the Scottish government published their NMP strategy, they were able to draw on a number of published papers providing evidence of the benefits [34], although in reality the only full independent prescribers included were nurses. Likewise the pharmacist prescriber training report in 2010 was also able to draw on practice examples to illustrate various different ways that independent prescribing had been implemented [35].

The 2010 White Paper ‘Equity and excellence: liberating the NHS’ signalled a change in direction for the health service, putting the patient at the centre of care with ‘*no decision about me without me*’ [75] but without the previous emphasis on workforce development; a point highlighted in a later staffing report [81]. The need for responsive and patient centred care, within the constraints of limited finances, was further developed in the subsequent Five-Year Forward View [16]. This document sets the need to provide more integrated care, giving patients greater control, against the background of increasing demand, rising costs resulting from new technologies, and budgetary constraints. Although prescribing is not specifically mentioned, there is a call to challenge traditional ways of working and to use the most appropriate healthcare professional for the task in hand.

This approach is echoed by the Welsh Assembly primary care plan, which describes a future model of primary-care in which the general practitioner acts as the leader over a multi professional team, who between them care for the patient [39]. The Welsh Assembly associated workforce development plan depends on other healthcare professionals taking on roles traditionally associated with general practitioners or secondary care, highlighting this with numerous examples of healthcare professionals taking on new or advanced roles [68]. One such example is the monitoring of low risk glaucoma patients by optometrists, and the document comments that there will be an increased need for optometrists to train as prescribers as they develop these advanced roles. NMP is perceived as integral to these developments. The English primary care report [40] describes a number of approaches to reducing the burden on general practitioners. Included in this are new models of practice such as the work of physicians’ associates, but as The Health Foundation comments, their role in relieving pressure on doctors will be limited if they are unable to prescribe [81]. The primary care report also describes prescribing in relation to physiotherapists, who are able to provide streamlined care for patients, and pharmacists to support their medicines optimisation activities, such as review of patients at risk of polypharmacy and adverse drug events [40]. Nurse prescribing is not specifically mentioned, although the report does identify that nurses have many responsibilities, including the care of patients with long-term conditions. More recently, the draft workforce strategy describes advanced practice for a number of professions such as nursing and paramedics but does not define what this entails [44]. It also describes podiatry and physiotherapy being potential first contact points for patients with musculoskeletal disorders. Prescribing would support all of these activities but is not explicitly mentioned and it could be perceived that NMP is seen to be so routine and embedded in practice for these professions that it warrants no mention. This compares with the pharmacy situation, where the same document put pharmacist independent prescribing as one of the priority areas to address and describes a project to put advanced pharmacists with prescribing skills into emergency departments. Other reports also make explicit mention of pharmacy prescribing as one of the tools to enhance medicines optimisation practices [41, 42] suggesting that pharmacist prescribing is still not embedded into routine practice.

A review of the professional distribution of policy documents supports this supposition concerning NMP becoming routine practice, with the majority involving generic NMP or covering multiple NMP professions (see **Error! Reference source not found.**). Of the three nursing specific policy documents, two date from before 2010, and the final one from 2017 [25, 28, 43]. Pharmacy alone of the professions is associated with multiple policy documents since 2013; with three by the Royal Pharmaceutical Society and one by each of the Scottish government and NHS England [36–38, 41, 42]. Similar recent policy documents were unable to be identified for any other of the NMP professions, despite in-depth searching. This may reflect the need for pharmacists to develop new roles and skills as the traditional dispensing role diminishes as a consequence of technological advances such as electronic prescribing and robotic dispensing. With medicines central to pharmacy practice, it is appropriate that these roles support medicines optimisation; however, these are not existing roles that a pharmacist can move into, rather they are roles that require creating. The pharmacy orientated policy documents are required to describe to both pharmacists and commissioners how pharmacist prescribing could work in practice. This compares with other healthcare professions, such as physiotherapy, where medicines formed an adjunct to their main practice area allowing roles to be expanded. Pharmacy could also be perceived to be an innately conservative profession, and the policy documents would thus serve to overcome a reluctance to adopt innovative working practices.

It is notable that there has been a shift regarding the role that NMP plays in the care of patients. The 2006 document, Medicines Matters, envisaged independent prescribers utilising a comparatively small personal formulary of drugs, which excluded controlled drugs and unlicensed medicines, to treat uncomplicated conditions [24]. Since independent prescribing for nurses and pharmacists was launched, their prescribing rights have been gradually extended to include unlicensed medicines and controlled drugs [46, 82] and more recent documents describe the role NMP has in the care of long-term conditions and complex patients [40, 42]. This is echoed by the changing role of medical staff in patient care. The early implementation guidance described medical staff retaining an overview of patient care, with nurse and pharmacist prescribing intended to improve patients access to medicines [23, 25, 26, 29]. Subsequent consultation processes (podiatry, physiotherapy, radiography and paramedics) have seen a change such that examples given in these documents describe the provision of a complete package of care without the need to involve other healthcare professionals. Indeed, the consequent reduction in costs through reducing appointments is listed as a benefit in the impact assessments [51, 56, 58, 61]. More recently, the Health Education England primary care report envisages that general practitioners will be treating patients with complex conditions, with other healthcare professionals providing routine care [40].

### Strengths and limitations

The strengths of the present review include the systematic, iterative approach to identifying relevant documents, using document mapping techniques to identify gaps in the evidence. The dynamic nature of this healthcare area inevitably means that this review provides a snapshot of the situation between 2006 and 2018, which may well be superseded by unanticipated developments. The selected papers relate to the UK and the devolved nations only and this may limit generalisability to other countries. Additionally, although the legislation permits the use of NMP in UK private healthcare, the policy documents concern the use of NMP in the NHS and this may further limit generalisability for alternative healthcare systems.

Despite extensive searches there may well be further policy documents available, such as from the home nations or professional bodies that are not identifiable through a search strategy.

## Conclusions

In conclusion it can be seen that this review has revealed that the government approach to NMP has changed over the 12 year period from 2006. Although originally intended as a means of improving patient choice and access to medicines, the emphasis has subtly changed to NMP supporting medical practitioners and reducing costs. Patients are expected to be cared for, and treated by, the most appropriate health care professional such as a physiotherapist for a musculoskeletal problem. This frees medical time, allowing medical practitioners to treat more complex cases. Costs are reduced by streamlining care through reducing multiple appointments with different healthcare professionals, and by using the most appropriately qualified professional.

This review has also highlighted the role that NMP now plays in patient care, with prescribing perceived as one skill in the advanced practice armamentarium used to treat and support patients, enabling patients to benefit from receiving a complete package of care from a single healthcare professional. As prescribing has become embedded into day to day practice for the majority of the NMP professions, so the need to highlight prescribing in policy documents has diminished (as seen in the recent workforce development document), just as it is no longer felt necessary to describe in detail advanced practice in these professions. As new models of practice are developed, such as use of physician’s associates, so the demand for NMP to expand to other healthcare professional groups continues, with the implication that prescribing is integral to these roles.

However, this review has found that while NMP has become embedded into routine practice for many professions, this is not universal. Despite pharmacists having achieved independent prescribing rights in 2006, it would appear from the repeated policy documents describing the need for pharmacist prescribers that it is still not embedded into pharmacists’ routine practice. Medicines remain at the core of pharmacy practice through supply and optimisation but, until the new roles become established, prescribing has yet to be perceived as a ‘normal’ pharmacist activity.

This review has also highlighted the impact that a change in government can have, as shown by the gap in policy document publication during the Coalition’s review and reorganisation of the NHS, and the delays in legislation concerning controlled drugs. These delays are not inevitable, as shown by the physiotherapist and podiatrist consultations which were conducted during this period.

While these findings concern a publicly funded health service in a single country, and may therefore be considered to have limited generalisability, there are messages that may resonate in other settings. These concern the impact of reorganisation on service development and how uptake of a novel skill is adopted by professions.

## Supporting information

S1 Appendix. PRISMA checklist

S2 Appendix. HMIC (Ovid) search strategy

S1 Protocol. PROSPERO record

